# Development of in-line anoxic small-angle X-ray scattering and structural characterization of an oxygen-sensing transcriptional regulator

**DOI:** 10.1101/2023.05.18.541370

**Authors:** Gabrielle Illava, Richard Gillilan, Nozomi Ando

## Abstract

Oxygen-sensitive metalloenzymes are responsible for many of the most fundamental biochemical processes in nature, from the reduction of di-nitrogen in nitrogenase to the biosynthesis of photosynthetic pigments. However, biophysical characterization of such proteins under anoxic conditions can be challenging, especially at non-cryogenic temperatures. In this study, we introduce the first in-line anoxic small-angle X-ray scattering (anSAXS) system at a major national synchrotron source, featuring both batch-mode and chromatography-mode capabilities. To demonstrate chromatography-coupled anSAXS, we investigated the oligomeric interconversions of the Fumarate and Nitrate Reduction (FNR) transcription factor, which is responsible for the transcriptional response to changing oxygen conditions in the facultative anaerobe *Escherichia coli*. Previous work has shown that FNR contains a labile [4Fe-4S] cluster that is degraded when oxygen is present, and that this change in cluster composition leads to the dissociation of the DNA-binding dimeric form. Using anSAXS, we provide the first direct structural evidence for the oxygen-induced dissociation of the *E. coli* FNR dimer and its correlation with cluster composition. We further demonstrate how complex FNR-DNA interactions can be studied by investigating the promoter region of the anaerobic ribonucleotide reductase genes, *nrdDG*, which contains tandem FNR binding sites. By coupling SEC-anSAXS with full spectrum UV-Vis analysis, we show that the [4Fe-4S] cluster-containing dimeric form of FNR can bind to both sites in the *nrdDG* promoter region. The development of in-line anSAXS greatly expands the toolbox available for the study of complex metalloproteins and provides a foundation for future expansions.

## Introduction

An estimated quarter to half of all proteins contain transition metal ions or metallocofactors.^1,2^ Notable biological processes catalyzed by metalloenzymes include the conversion of dinitrogen to ammonia by nitrogenases, oxygen evolution by photosystem II, H_2_ production by hydrogenases, and carbon cycling by carbon monoxide dehydrogenases.^3–6^ Structural characterization of such proteins often require special strategies to maintain the integrity of the metallocofactor, which can be sensitive to a number of environmental factors.^7^ In particular, molecular oxygen can readily accept electrons from transition metals.^8,9^ Such interactions can lead to unwanted changes in the oxidation state of the metallocofactor as well as lead to destabilization^9^ or degradation of the protein scaffold.^10^ In crystallography and cryo-electron microscopy, oxygen-sensitive samples can be frozen within an anoxic environment and maintained at cryogenic temperatures during data collection.^7,11,12^ Alternatively, serial femtosecond crystallography may be pursued for room-temperature data collection.^13^ However, solution-based analyses, such as small-angle X-ray scattering (SAXS), require special experimental accommodations.

SAXS, although a relatively low-resolution technique, is a powerful way to characterize protein-protein interactions and conformational changes under near physiological conditions.^14^ Recent advances in chromatography-coupled SAXS and advanced data processing algorithms, such as evolving factor analysis (EFA)^15^ and regularized alternating least squares (REGALS)^16^, have opened the way for analysis of samples which exist in solution as a mixture of states. However, such modern SAXS analyses have not been generally available for oxygen-sensitive proteins. Traditionally, stringent anoxic conditions were maintained by loading and sealing samples in thin-walled glass capillaries inside of an anoxic chamber and then transporting these samples to an X-ray source for static data collection.^17^ Although such an approach ensures anoxic conditions for long periods, it is problematic for a multitude of reasons. Capillary scattering is difficult to reproduce and subtract exactly. Moreover, this approach is not amenable to biological samples that are not stable for long periods of time, nor does it allow for sample flow within the X-ray beam, which is commonly used to minimize radiation damage and to separate mixtures via in-line chromatography. To expand the experimental capabilities available for the study of oxygen-sensitive metalloproteins, many of which are structurally complex, it is necessary to design and implement a system in which advanced SAXS experiments can be performed in the absence of oxygen.

An ideal test case for the development of anoxic SAXS is the Fumarate and Nitrate Reduction transcription factor (FNR) from *Escherichia coli*.^18^ FNR is a member of the cyclic adenosine monophosphate (cAMP) receptor protein (CRP)/FNR superfamily of transcriptional regulators. Members of this family are known for three structural features: an N-terminal effector-binding domain, a C-terminal DNA-binding domain, and a dimerization interface.^19^ Many members of this family are homodimers that when bound to specific effectors, can then bind to DNA to regulate transcription. Upon effector binding, a structural change is propagated through the protein to the DNA-binding region, thus making this family a compelling target for structural analysis.^19,20^ The effector-binding region of *E. coli* FNR is unusual in that it contains a [4Fe-4S] cluster that initially degrades to a [2Fe-2S] cluster upon oxygen exposure, which ultimately converts to the apo, cluster-less form *in vivo*.^21–24^ The local amino acid rearrangement caused by the degradation of the [4Fe-4S] cluster is thought to propagate to the dimerization interface and lead to dissociation of the homodimer, which in turn prevents FNR from binding to DNA.^18,25^ In this way, FNR acts as an oxygen sensor within facultative anaerobes, reacting to changes in oxygen availability through FeS cluster state and acting as a global transcriptional regulator to facilitate the transition between aerobic and anaerobic metabolisms. FNR function has been extensively studied in the facultative anaerobe *E. coli* through mutational analysis and *in vivo* fitness of cells, in addition to spectroscopic characterization of metal cluster content as a function of oligomerization state.^18,21,26–30^ However, direct structural characterization of *E. coli* FNR has been challenging.^31^

In this study, we describe the first in-line, anoxic SAXS (anSAXS) experimental set-up that is open to all researchers as a technique at a major national synchrotron. This system allows for either chromatography-coupled SAXS or batch-mode SAXS to be performed in the absence of oxygen without relying on chemical reductants. Additionally, the system provides the capability to measure UV-Vis absorption spectra during sample elution in chromatography mode, which enables monitoring of metallocofactors with distinct spectral characteristics that can serve as an additional source of information. Utilizing chromatography-coupled anSAXS, we demonstrate the absence of oxygen contamination through analysis of dimeric *E. coli* FNR with an intact [4Fe-4S] cluster. We further show that briefly exposing this sample to air leads to a monomeric form of *E. coli* FNR with a [2Fe-2S] cluster. Finally, we combine UV-Vis analysis and chromatography-coupled anSAXS to characterize *E. coli* FNR interactions with a native promoter sequence that controls the transcription of the anaerobic ribonucleotide reductase genes (*nrdDG*), which enable *de novo* production of nucleotide precursors for DNA biosynthesis and repair in the absence of oxygen. The introduction of in-line anSAXS greatly expands the toolbox available to the study of oxygen-sensitive metalloproteins and provides opportunities for future expansions, such as time-resolved and high-pressure anSAXS.

## Results

### Design of in-line anoxic SAXS setup

To enable in-line SAXS in the absence of oxygen, we designed a dual-mode system within an anoxic chamber, which interfaces with an existing in-vacuum X-ray flow cell at beamline ID7A of the Cornell High Energy Synchrotron Source (CHESS)^14^ (Figure 1). The anoxic chamber (Type C, Coy Lab Products, Grass Lake, MI) houses two sample ports: an injection port (Rheodyne MXX777-601, IDEX Health & Science, Rochester, NY) for chromatography-coupled anSAXS and a sample funnel for pipette-based loading into the X-ray flow cell. For chromatography mode, the flow rate and pressure limits are controlled by an external pump (LC-20AD, Shimadzu North America, Columbia, MD) (Figure 1, pink path). In batch-mode experiments, a syringe pump (Cavro XCalibur Modular Digital Pump, Tecan Systems, San Jose, CA) inside the chamber is used to control the delivery and oscillation of direct injection samples (Figure 1, blue path). Conventional 10-32 coned 1/16” FPLC fittings and polyetheretherketone (PEEK, 0.25 mm ID chromatography mode, 0.76 mm ID batch mode) tubing are used to construct the flow paths and to connect to the external X-ray cell. The flow path is fully closed, such that the sample is returned into the chamber after X-ray exposure. In chromatography mode, the sample then flows through an in-line UV-Vis cell (SMA-Z Cell 79048, 2.5 mm optical path, FIAlab, Seattle, WA) with a fiber-optic coupled spectrometer (AvaSpec-ULS2048, Avantes, NS Apeldoorn, The Netherlands). A halogen-deuterium light source (Avalight-DH-S, Avantes, NS Apeldoorn, The Netherlands) outside of the chamber provides 200-800 nm light for full spectral analysis of biological samples. Both the light source and the Shimadzu pump are kept outside of the anoxic chamber to reduce heating within the chamber. With this setup, we are also able to house other necessary tools within the chamber, such as a refrigerated centrifuge for sample preparation and a nanophotometer for sample characterization.

**Figure 1:**
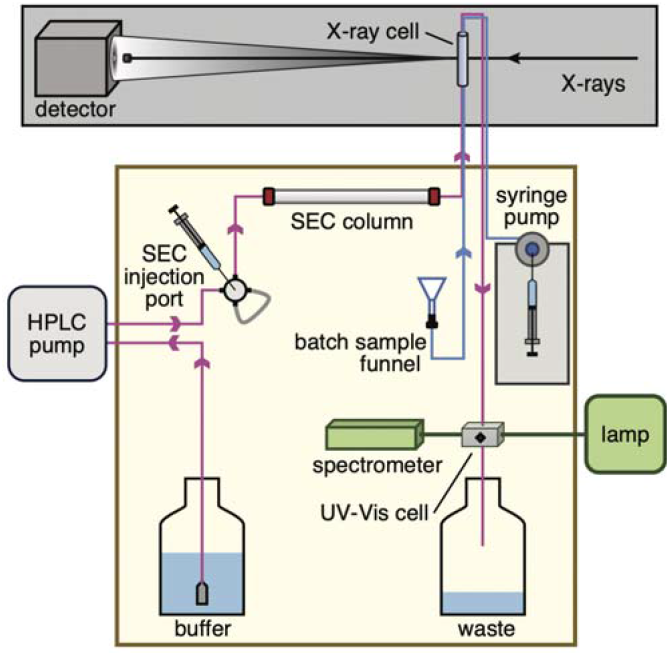
Schematic of anoxic SAXS system. An anoxic atmosphere is maintained by a vinyl glove box (khaki box; gloves and airlock not shown) in the X-ray hutch at beamline ID7A at the Cornell High Energy Synchrotron Source. The glove box contains two sample-loading paths: in-line chromatography (pink) and batch-mode (blue). It is a closed system, with all tubing openings contained within the anoxic environment. In chromatography mode, buffer is pulled through an external high-performance liquid chromatography (HPLC) pump, then used to propel injected sample through a size exclusion chromatography (SEC) column, X-ray sample cell, and UV-Vis sample cell. The fiber-coupled spectrometer and external UV-Vis light-source enable time-dependent UV-Vis absorption measurements over the wavelength range 200-800 nm. In batch mode, 30-50 μL sample is loaded into the batch sample funnel, and a syringe pump controls its flow and oscillation during data collection. The X-ray cell and detector are within a vacuum flight path.

This dual-mode system has several useful features. The in-line spectrometer produces a full UV-Vis spectrum, which is particularly useful for metal-containing proteins that have unique spectroscopic signatures. A time series can be measured at any wavelength and aligned to a SAXS series, so the sample elution can be characterized by both X-ray scattering and UV-Vis absorption during the same experiment. Measuring the absorption of the sample post X-ray exposure is also advantageous to ascertain if there has been a change in the oxidation state of a metallocofactor as a function of radiation. Furthermore, absorption spectra can be used to verify the absence of contaminating oxygen, an important validation step after the sample travels outside of the chamber through the X-ray sample cell tubing. To provide experimental capabilities comparable to conventional SAXS at ID-7A, we installed batch-mode components to enable manual pipette-based loading of individual 30-50 μL samples. This mode is essential for data collection on titration series, concentration series, or time-sensitive samples which require immediate data collection. This mode uses the syringe pump to pull the sample through the tubing into the X-ray sample cell and oscillate the sample plug in the X-ray beam to reduce radiation damage. Between samples, cleaning solutions are introduced into the X-ray cell via this mode, followed by rinsing with 100% methanol and drying with N_2_ gas, which is fed into the chamber through a valved port. This batch mode is nearly analogous to conventional aerobic SAXS at ID-7A and utilizes the same user interface to control sample movement.

Switching from conventional SAXS to anSAXS can be done rapidly by wheeling the anSAXS system next to the X-ray path and connecting the tubing between the chamber and the X-ray sample cell. The anoxic chamber is sealed and deoxygenated for at least four days prior to the first experiments of a scheduled run. The oxygen level within the chamber is monitored through an O_2_/H_2_ sensor (CAM-12, Coy Lab Products) with accuracy between 20-700 ppm O_2_ and 1-10% H_2_. For our anSAXS system, the H_2_ concentration is maintained at >2% for the efficiency of palladium catalyst packs used to scavenge O_2_. Catalyst packs are regenerated within a vacuum oven and replaced every 24-48 hrs of an experiment. All plasticware that encounters the sample during an experiment, including the tubing and consumables (e.g., sample tubes and pipette tips), are degassed by a 72-hr incubation in a vacuum oven at 100°C, followed by a 24-hr treatment under the antechamber vacuum in order to be sufficiently free of oxygen. Prior to use, all buffers and cleaning solutions are filtered, sparged with nitrogen, and allowed to equilibrate in the anoxic chamber through vigorous stirring overnight. Prior to each experiment, 20 mM sodium dithionite solution is flushed through the flow path to scavenge any contaminating oxygen from the surface of the tubing, followed by deoxygenated deionized water. The absence of contaminating oxygen is confirmed by measuring the UV-Vis absorbance of indigo carmine, a redox sensitive indicator dye, that is left open to the chamber atmosphere, as well as upon mixing with flow-through of deionized water that has passed through the tubing.

To transport stock solutions and deoxygenated buffers to the beamline, we recommend shipping them frozen on dry ice or unfrozen in crimped vials, depending on the sample requirements. For short periods (e.g., 20 min), it is possible to transport liquids within glass flasks with airtight valves or in gas-tight transport containers containing catalyst packs (BD GasPak™ EZ, Anaerobe Container System, 260678-20). Items brought into the anoxic chamber undergo two antechamber cycles, and the O_2_ level within the chamber is maintained at <30 ppm as measured by the O_2_/H_2_ sensor. When changing chromatography columns, a minimum of 5 column volumes (CVs) of deoxygenated buffer is passed through the column to ensure that it is anoxic, with the first 2 CVs containing at least 10 mM sodium dithionite. When changing buffer within an already deoxygenated column, 2 CVs of buffer are passed through. Column equilibration can coincide with batch-mode experiments because each experimental method has a separate and dedicated flow path.

### Oxygen dependence of the FNR quaternary structure in solution

To demonstrate the performance of our anSAXS setup, we chose to examine FNR, an oxygen-sensing protein. Previously, it was shown with size-exclusion chromatography (SEC) that FNR undergoes a change in the oligomerization state that is correlated with the assembly and degradation of its oxygen-sensitive [4Fe-4S] cluster.^18,26,30^ For direct comparison with prior work, we investigated *E. coli* FNR using size-exclusion chromatography coupled with anSAXS (SEC-anSAXS).

*E. coli* FNR with an Fe content of 3.27 ± 0.48 per monomer was prepared anaerobically at 8.5 mg/ml (328 μM monomer concentration) in FNR buffer (100 mM Tris HCl pH 6.8, 150 mM NaCl, 10 % glycerol). A 100-μL aliquot, which was kept anoxic, was injected onto a deoxygenated and pre-equilibrated Superdex 75 Increase 10/300 column (Cytiva Life Sciences, Marlborough, MA) at a flow rate of 0.5 ml/min. Scattering was measured continuously during the elution, and scattering profiles prior to the protein elution were used to subtract the background scattering from the full dataset. The resulting SEC-anSAXS dataset is plotted as *I* vs *q* vs frame number (Figure S1A, left), where *I* is the scattering intensity, and the momentum transfer variable is defined as *q* = 4π sin(θ)/λ. Here, 2*θ* is the scattering angle, and λ is the X-ray wavelength. This experiment was repeated with a second 100-μL aliquot from the same sample preparation, which was exposed to air for 5 min prior to data collection (Figure S1B, left). The Fe content of the oxygen-exposed sample was 1.91 ± 0.48 per monomer.

The dataset collected on the anoxic sample shows a dominant peak with two shoulders on either side (Figure 2A, top black curve and Figure S1A, middle). Consistent with the radius of gyration (*R*_*g*_) changing across the elution (Figure S1A, middle), singular value decomposition (SVD) indicated that the dataset consisted of 4 significant components (Figure S1A, right).^15^ Using REGALS, we were able to decompose the dataset into 3 overlapping protein peaks (Figure 2A, top) and one changing background component (Figure S2).^16^ The dominant protein component (Figure 2A, top blue peak) has an *R*_*g*_ value of 26.0 ± 0.1 Å by Guinier analysis, which agrees well with the theoretical value of 25.8 Å, calculated from a model of the *E. coli* FNR dimer derived from AlphaFold2.^32^ The two minor components correspond to a larger species (Figure 2A, top red peak) with an *R*_*g*_ value of 34.8 ± 0.5 Å and a smaller species (Figure 2A, top orange peak) with an *R*_*g*_ value of 22.2 ± 0.2 Å. The latter *R*_*g*_ value is similar to the theoretical value calculated from an AlphaFold2 model of the *E. coli* FNR monomer (21.5 Å). Similar analyses were performed for the dataset collected on the oxygen-exposed sample (Figures S1B). REGALS was used to decompose the dataset into 2 slightly overlapping protein peaks (Figure 2A, bottom) and a changing background component (Figure S3). For this sample, the first, minor component (Figure 2A, bottom blue peak) has an *R*_*g*_ value suggestive of a dimer (25.4 ± 0.6 Å), while the dominant protein component (Figure 2A, bottom orange peak) has an *R*_*g*_ value suggestive of a monomer (21.3 ± 0.1 Å).

**Figure 2:**
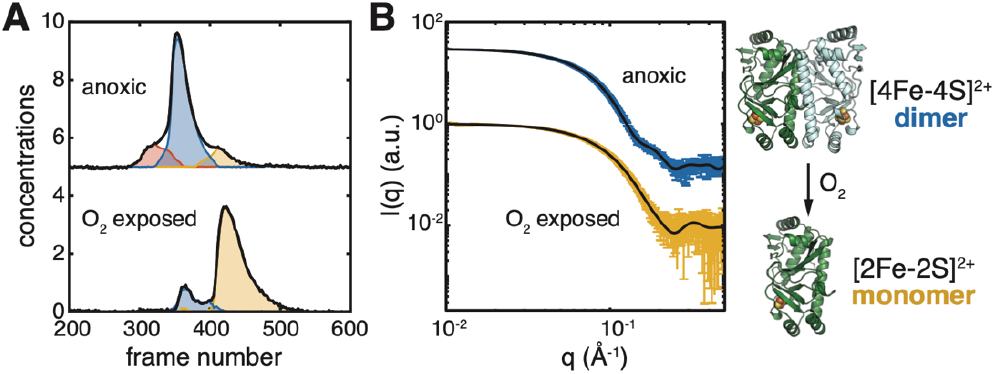
SEC-coupled SAXS analysis of anoxic and O_2_-exposed *E. coli* FNR. All samples were in 100 mM Tris HCl pH 6.8, 150 mM NaCl, 10 % glycerol; FNR concentrations are dimer concentrations. **(A)** Top: SEC-SAXS elution profile of 164 μM *E. coli* FNR (black) with an Fe content 3.27 ± 0.48 per monomer that kept was anoxic. This dataset was decomposed into three protein components (red, blue, orange) by REGALS analysis (Figure S2), having *R*_*g*_ values of 34.8 ± 0.5, 26.0 ± 0.1, and 22.2 ± 0.2 Å. Bottom: SEC-SAXS elution profile of an aliquot of the sample described in panel A that was exposed to air for 5 min prior to data collection (black), which led to an Fe content of 1.91 ± 0.48 per monomer. This dataset was decomposed into two protein components (blue, orange) by REGALS analysis (Figure S3), having *R*_*g*_ values of 25.4 ± 0.6 and 21.3 ± 0.1 Å. **(B)** The scattering profile of the dominant species in the anoxic dataset (blue) is well-described by the theoretical scattering (black) of the AlphaFold2 model for the *E. coli* FNR dimer (shown to the right) with a χ^2^ of 1.58. The scattering profile of the dominant species in the oxygen-exposed dataset (orange) is well-described by the theoretical scattering (black) of the AlphaFold2 model for the *E. coli* FNR monomer (shown to the right) with a χ^2^ of 1.08. Altogether, these results indicate that oxygen exposure leads to the dissociation of [4Fe-4S] cluster-containing dimer into [2Fe-2S] cluster-containing monomer.

Structural modeling was then performed on scattering profiles of the dominant species in each dataset. We find that the scattering profile from the dominant species in the anoxic sample (Figure 2A, top blue peak) shows excellent agreement with the theoretical scattering calculated from the dimer model (Figure 2B, blue and black), while that of the oxygen-exposed sample (Figure 2A, bottom orange peak) shows excellent agreement with that of the monomeric model (Figure 2B, orange and black). Overall, these results demonstrate that SEC-anSAXS can be performed without oxygen contamination and in the absence of reductant. Together with Fe content analysis, they provide the first structural evidence in support of the allosteric model for *E. coli* FNR, in which a dimer with a [4Fe-4S] cluster degrades to a monomer with a [2Fe-2S] cluster upon exposure to oxygen.

### FNR-DNA interactions under anoxic conditions

To fully test the capabilities of SEC-coupled anSAXS, we examined the interactions of *E. coli* FNR with a native DNA sequence. One of the key changes in the transition from aerobic to anaerobic metabolism in *E. coli* is the mechanism by which deoxyribonucleotides are produced for DNA synthesis. Under aerobic conditions, *E. coli* uses a class I ribonucleotide reductase (RNR) to convert ribonucleotides to deoxyribonucleotides and switches to class III RNR usage under anaerobic conditions.^33–35^ Unlike class I RNRs, which require oxygen to assemble a catalytically essential cofactor, class III RNRs use Fe-S and glycyl radical chemistry and are therefore highly oxygen sensitive.^36^ Previously, it was shown by gel-shift assays that FNR controls the transcription of the class III RNR genes (*nrdDG*) in *E. coli* by binding the promoter region, which contains two FNR consensus motifs: one centered at the -33.5 position from the transcription start site (site-1) and another centered at the -64.5 position (site-2).^37^ To test this observation, we performed a binding analysis of *E. coli* FNR with the *nrdDG* promoter sequence using SEC-anSAXS.

Anaerobically prepared *E. coli* FNR with an Fe content of 2.43 ± 0.72 per monomer was mixed with double-stranded 54-bp oligonucleotide containing the *nrdDG* promoter sequence and concentrated to account for the nearly 10-fold dilution on the SEC column. The final mixture, which contained 100 μL of 360 μM *E. coli* FNR (dimer concentration) and 160 μM oligonucleotide, was then injected onto a deoxygenated and pre-equilibrated Superdex 75 Increase 10/300 column and eluted with FNR buffer at a flow rate of 0.5 ml/min. The SEC-anSAXS dataset collected on the FNR-DNA sample shows a dominant peak followed by two smaller peaks (Figure 3A, top black curve and Figure S1C). SVD of the entire dataset produces four significant components (Figure S1C, right), with the central region of each peak contributing one significant value. Based on this result, REGALS analysis was performed to decompose the dataset into 3 overlapping protein peaks and a changing background (Figure 3A, top and Figure S4).

**Figure 3.**
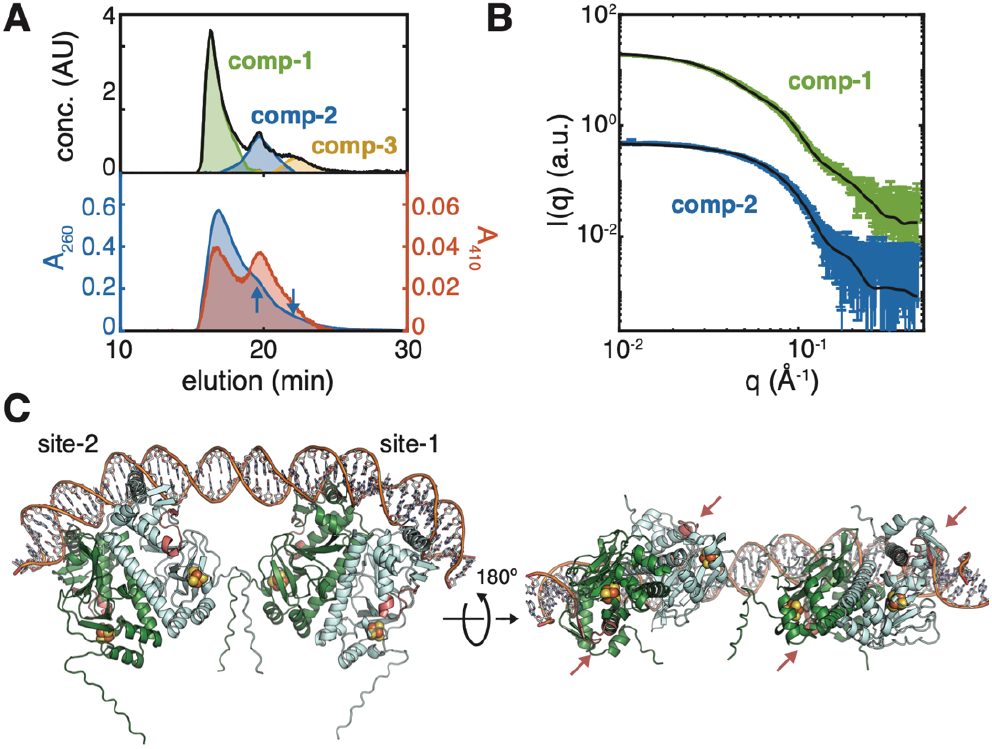
SEC-coupled SAXS analysis of *E. coli* FNR bound to the *E. coli nrdDG* promoter sequence. **(A)** Top: SEC-SAXS elution profile (black) of 360 μM *E. coli* FNR with 160 μM oligonucleotide containing the *E. coli nrdDG* promoter sequence in 100 mM Tris HCl pH 6.8, 150 mM NaCl, 10% glycerol. This dataset was decomposed into three protein components (green, blue, orange) by REGALS analysis (Figure S4), having *R*_*g*_ values of 44.5 ± 0.2, 28.1 ± 0.2, and 23.3 ± 0.3 Å. Bottom: Absorbance at 260 nm (blue) correlates strongly with component 1 (top, green), which elutes at 17.0 min, suggesting that this species contains DNA. Small shoulders at 19.7 min and 22.1 min (blue arrows) correlating with components 2 and 3 likely arises from protein absorbance. Absorbance at 410 nm correlates with both components 1 and 2 (green and blue in top panel), suggesting that these species contain FeS clusters. **(B)** The scattering profile extracted from REGALS analysis for component 1 (green) is well-described by the theoretical scattering (black, χ^2^ = 2.48) of a model of FNR doubly bound to DNA (shown in panel C), while that of component 2 (blue) is described by the theoretical scattering (black, χ^2^ = 1.75) of the Alphafold2 model for the *E. coli* FNR dimer shown in Figure 2. **(C)** Model of FNR doubly bound to the *E. coli nrdDG* promoter sequence. Site-1 and site-2 are centered at -33.5 and -64.5 bp from the transcription start site. FNR interacts with RNA polymerase via the activation region 1 (AR1), which is shown in salmon (indicated by arrows in right panel). The FNR N-termini (pointing downwards from FNR in the left panel) are disordered in this model.

The absorbance was simultaneously measured at 410 nm and 260 nm (Figure 3A, bottom) to monitor the [4Fe-4S] cluster and DNA contents, respectively. The 410 nm trace contains two prominent peaks at 16.9 and 19.8 min (Figure 3A, bottom red) that coincide with the first two eluting components (Figure 3A, top green/blue peaks), suggesting that these species contain an intact [4Fe-4S] cluster. The 260 nm trace, on the other hand, starts with a dominant peak at 17.0 min followed by two small shoulders at 19.7 and 22.1 min (Figure 3A, bottom blue) that correlate well with each of the three components observed in the SEC-anSAXS elution (Figure 3A, top). The strong A260 absorbance signal at 17.0 min suggests that the first eluting species contains both protein and DNA, while the much weaker signal in the following peaks (Figure 3A, bottom blue arrows) likely arises from protein absorbance alone and suggest that the second and third species do not have bound DNA.

Guinier plots for the three REGALS-separated profiles are shown in Figure S4D. The *R*_*g*_ for the second species is 28.1 ± 0.2 Å, in excellent agreement with the experimentally derived value for the FNR dimer. Likewise, the *R*_*g*_ for the third species (23.3 ± 0.3 Å) is in reasonable agreement with the experimentally derived value for the FNR monomer. Combined with the FeS cluster absorbance monitored at 410 nm, these *R*_*g*_ values indicate that the second eluting species is the free FNR dimer with an intact [4Fe-4S] cluster and that the third species is a monomer lacking an intact cluster. The presence of apo-FNR is not unexpected as the overall Fe content of the sample was less than 4 per monomer. In contrast, the *R*_*g*_ of the first eluting species is significantly larger with a value of 44.5 ± 0.2 Å, which when combined with the strong 260 nm signal, is indicative of protein-DNA complex formation.

To determine the overall shape of the FNR-DNA complex, we next compared the pair distance distribution functions, *P*(*r*), of the three dominant species characterized in this study (Figure S5). *P*(*r*) provides shape information on a pure sample in real space and can be thought of as a histogram of intraparticle distances, *r*. As expected, dimerization of FNR leads to an increase in maximum dimension, *D*_*max*_ (Figure S5, orange to blue), but the overall shape of the *P*(*r*) curves remains monomodal, indicating that both the monomer and dimer are largely globular. However, the *P*(*r*) curve of the FNR-DNA complex exhibits a highly skewed shape (Figure S5, green) with a very long dimension (*D*_*max*_ ∼170 Å), indicating that it is overall an elongated species. Notably, there is a peak at small *r* (close to the peak position of the FNR dimer) and a shoulder at large *r* (Figure S5, green). This shape suggests that the FNR-DNA complex likely consists of two FNR dimers that are held apart like a dumbbell.

To create a model for two FNR dimers bound to DNA, we first examined all protein-DNA contacts in the crystal structure of a homologous protein, *Bradyrhizobium japonicum* FixK_2_, bound to a 30-bp double stranded oligonucleotide (PBD: 4i2o).^38^ In this structure, the helix-turn-helix motifs in the FixK_2_ dimer are bound to a pseudosymmetric pair of the canonical FNR consensus sequence (TT<u>G</u>ATnnnnA<u>TC</u>AA) (Figure S6A). It was previously shown that Arg200 and Glu196 in each monomer of FixK_2_ is directly involved in hydrogen bonding with the bases of a G·C pair and an adjacent T (underlined in the FNR consensus sequence) (Figure S6B). These residues and four other residues involved in interacting with the DNA phosphate backbone are fully conserved in *E. coli* FNR. Based on these observations, we aligned the AlphaFold2 model of *E. coli* FNR with the FixK_2_ crystal structure and mutated the DNA sequence to create models of FNR bound to each binding site in the *nrdDG* promoter sequence (Figure S7). The two sites were then connected by B-form DNA with the intervening sequence, and geometry optimization was performed after confirming that all base pairs were correctly oriented. The final model (Figure 3C) preserves the previously observed DNA curvature in each binding site and has a theoretical *R*_*g*_ of 47.1 Å. The theoretical scattering from this model agrees remarkably very well with experimental profile with no further refinement (Figure 3B, green and black). The agreement of this model with both the scattering data and the UV-Vis absorbance data suggests that both FNR sites on the *nrdDG* promoter region are occupied under the conditions of this experiment.

## Discussion

Oxygen-sensitive metalloenzymes are responsible for many of the most fundamental biological reactions in all kingdoms of life. Such compelling examples motivate the development of new capabilities that enable the characterization of these biomolecules in as close to physiological condition as possible. SAXS is a powerful tool in this respect, providing insight into protein-protein interactions and structural changes as a function of solution condition (e.g., temperature, pressure, pH, salt, ligand concentration) and other parameters (e.g., time). Here, we introduced an in-line system at CHESS beamline ID7A that enables both chromatography-coupled SAXS and batch-mode SAXS under anoxic conditions. The effectiveness of batch-mode anSAXS was recently demonstrated by an investigation of the cobalamin-dependent methionine synthase, in which the highly oxygen-sensitive Cob(I) oxidation state of the enzyme was maintained throughout data collection.^39^ In this current study, we report the first demonstration of SEC-anSAXS through analysis of the oxygen-dependent oligomerization and DNA binding of *E. coli* FNR. Because *E. coli* FNR can exist as a mixture of oligomeric species, the introduction of in-line chromatography was an essential development. By coupling SEC-anSAXS with full UV-Vis absorption spectra, we were able to further support the identities of the FNR-DNA complex and [4Fe-4S] cluster-containing FNR dimer.

Despite several decades of intensive study, structural characterization of *E. coli* FNR has been challenging, and the long sought-after crystal structure has remained elusive.^31^ It has been shown previously through mutagenesis, SEC, and oxygen exposure tests, that the active DNA-binding form of *E. coli* FNR is dimeric and contains an oxygen-labile FeS cluster.^18,22,23,25,26,30^ We have expanded on previous multi-wavelength SEC analysis^26^ through orthogonal SAXS data collection that definitively shows the connection between the intact [4Fe-4S] cluster and the dimeric state. Furthermore, by applying advanced deconvolution techniques,^15,16^ we were able to extract pure scattering profiles for each species within overlapping elution peaks and perform model-fitting. In doing so, we found that the change in oligomerization state can be described largely by rigid-body models of the FNR dimer and monomer, suggesting that large-scale change in tertiary structure are not required, as was suggested in previous proposals.^22^ Rather, our data support the hypothesis that shifts in the residues immediately surrounding the FeS cluster are subtly translated through the protein to the dimerization interface.^26,28^

Further, we extended this analysis to include the endogenous *E. coli nrdDG* promoter region that is known to contain two canonical FNR consensus motifs.^37,40^ Previous work by Roca *et al*.^37^ showed differential binding affinities for the two sites on the *nrdDG* promoter region using an oxygen-tolerant D154A mutant^30^ of FNR (FNR*). Using electrophoretic mobility shift assays, it was shown that the FNR* has a higher binding affinity to site-2 than site-1. This is not surprising as site-1 lacks one of the canonical CT/G triads in the consensus sequence (Figure S7A). By mutating the FNR consensus motifs, it was further shown that FNR* binding to site-2 is necessary and sufficient for initiating expression of the *nrdDG* gene, while binding site-1 only increases expression when site-2 is occupied.^37^ This dynamic would allow a modulated physiological response to decreasing levels of oxygen. In our study, we were able to perform the first structural characterization of DNA binding by wild-type *E. coli* FNR under anoxic conditions. As the SEC column leads to a nearly 10-fold dilution, we performed our experiment at high concentrations of FNR and DNA to observe maximal complex formation. Under these conditions, we did not observe a singly bound FNR-DNA complex. Using SVD and REGALS analyses, we showed that the SEC-anSAXS dataset can be described by three eluting species that are well-described by a doubly bound FNR-DNA complex, FNR dimer, and FNR monomer. Moreover, none of the species agree with a model of a singly bound FNR-DNA complex (fitting such a model yields χ^2^∼9 and 20 for components 1 and 2, respectively). In our model of the doubly bound FNR-DNA complex, the two FNR dimers are bound on the same side of the DNA. The curvature of the DNA brings the two dimers in close proximity at their N-termini (Figure 3C). If there are any interactions between the two FNR dimers that could lead to cooperative binding, we would propose that the flexible N-termini would need to be involved. A remaining mystery is how FNR binding at these two sites on the *nrdDG* promoter region leads to the activation of RNA polymerases. In our model, the so-called activation region 1 (AR1) of FNR, which is known to interact with class I and II RNA polymerases^41^, points away from the DNA axis at an angle (Figure 3C, salmon loops indicated by arrows). Now that a workflow for study FNR-DNA complexes has been established with SEC-coupled anSAXS, future studies can be aimed at elucidating the structure(s) of FNR in complex with RNA polymerase(s) and DNA.

At the writing of this text, fully anoxic in-line SAXS capability of this type has not yet been established at any other major national synchrotron. Here, we have shown the utility of SEC-coupled SAXS in obtaining cleanly separated scattering signals from heterogenous samples, and the value of full-spectrum UV-Vis traces in confirming the identities of the different sample states in solution. Moving forward, we plan to integrate the UV-Vis analysis into the existing software used at the beamline for SAXS data analysis, BioXTAS RAW^42^, and further simplify the batch-mode protocol by developing automated cleaning. More importantly, the in-line anSAXS system can accommodate new experimental modes. Currently, the development of anoxic high-pressure SAXS is underway. Such capabilities would enable to the study of redox-active proteins and metalloenzymes from deep sea organisms and other extremophiles. Additionally, time-resolved anSAXS is an exciting area to pursue in future work. Indeed, the outlook for anoxic experimental capabilities, is bright.

### Experimental Procedures

#### Protein Expression and Purification

An *E. coli* strain that is a Δ*fnr*, Δ*crp* BL21(DE3) derivative (PK22) containing plasmid encoding for wild-type FNR (pPK823)^43^ was kindly provided by Prof. Patricia Kiley (Univ. Wisconsin Madison). FNR was expressed and purified following established protocols^44^ with several modifications. Glycerol stocks (30% (v/v)) were prepared from individual colonies. The starter culture was prepared by inoculating 150-200 mL of LB media supplemented with 0.2% glucose and 100 mg/L ampicillin and overnight shaking at 200 rpm at 37 °C. Large-scale cultures were prepared by adding 15-20 mL of saturated starter culture (volume adjusted to reach an initial OD_590_ of ∼0.05) into each 1 L flask of temperature-equilibrated M9 minimal media supplemented with 0.2% glucose, 0.2% casamino acids, 4 mg/L thiamine, 1 mM MgSO_4_, 1 mM CaCl_2_, 10 mg/mL ferric ammonium citrate, and 100 μg/mL ampicillin, as described previously.^44^ The large-scale cultures were incubated at 37 °C, 180-220 rpm to an OD_590_ of 0.3-0.5, at which point isopropylthio-β-galactoside (IPTG) was added to a final concentration of 0.4 mM. After 2 hours of induction, ferric ammonium citrate was added to a final concentration of 20 mg/L before the cultures were transferred to a cold room. After continuous sparging with nitrogen gas for 14-16 hours at 4 °C, cells were harvested by centrifugation at 3,500 rcf for 20 minutes at 4 °C. Cell paste was either flash frozen in liquid nitrogen for storage or immediately transferred to an anaerobic COY chamber (96% N_2_, 4% H_2_, <30 ppm O_2_ atmosphere) for cell lysis. Chemical lysis was performed by resuspending cells in 5 mL/g cell paste deoxygenated lysis buffer (100 mM Tris HCl pH 6.8, 75 mM NaCl, 10% glycerol) supplemented with 0.5 mg/mL hen egg white lysozyme (Sigma, L6876-1G, #SLBT5161), 5 units/mL Benzonase nuclease (EMD Millipore, 707463, #377108), 20 μM phenylmethylsulfonyl fluoride in ethanol (PMSF), 1.5 mM MgCl_2_, 0.5% (v/v) Triton X, 2 mM sodium dithionite. All plasticware in contact with protein was degassed in vacuum under heat for 72 hours and equilibrated at room temperature in an anaerobic atmosphere for at least 24 hours prior to use.

All purification steps were performed anaerobically in the COY chamber using an ÄKTA go FPLC system (Cytiva). Post lysis, cell debris was separated from clarified lysate by centrifugation at 12,000 rcf for 30 minutes at 4 °C. Lysate was sequentially passed through 0.7-μm and 0.2-μm syringe filters prior to loading onto a 10 mL HiTrap SPFF anion exchange column pre-equilibrated with 5 CVs of deoxygenated wash buffer (100 mM Tris HCl pH 6.8, 75 mM NaCl, 10% glycerol). The loaded column was washed with 10 CVs of the wash buffer before eluting with a 50-mL linear gradient from 0.075 to 1 M NaCl at a flow rate of 2 mL/min. Fractions containing FNR were pooled, concentrated, and loaded onto a HiLoad 16/600 Superdex 75 pg column preequilibrated with 100 mM Tris HCl pH 6.8, 150 mM NaCl, 10% glycerol to separate FNR dimer, FNR monomer, and lysozyme. The FNR concentration was determined by Bradford assay^45^ using bovine serum albumen (Research Products International Corp., 9048-46-8, #33403) as a standard, and the Fe content was determined by Ferrozine assay^46^. Errors associated with Fe content are standard deviations of the [Fe]/[FNR] ratio, calculated from the standard deviations of individual [FNR] and [Fe] measurements, assuming that they are independent of each other.

### DNA Preparation

Complementary 54-mer oligonucleotides were purchased from Integrated DNA Technologies to obtain a double-stranded DNA fragment containing the sequence for the *E. coli* class III RNR (*nrdDG*) promoter region^37,47^ (5′-TACT<u>TTGAG</u>CTAC<u>ATCAA</u>AAAAAGCTCAAACATCC<u>TTGAT</u>GCAA<u>AGCAC</u>TATAT-3′, where the FNR consensus sequence is underlined). Lyophilized single-stranded DNA was resuspended in 50 mM Tris HCl pH 8.0, 75 mM NaCl, 5% glycerol, 2 mM sodium dithionite within the COY chamber. No EDTA was added as nuclease inhibitor since it interacts unfavorably with Fe-S clusters, however, RNAse and DNAse free plasticware was used for all solutions containing DNA.^48^ DNA concentration was monitored by absorbance at 260 nm. Single-stranded oligonucleotides were stored at 100 μM and diluted to 10 μM for annealing. Degassed 96-well PCR compatible plates and covers were filled with 104 μL/well 10 μM of both complementary single stranded oligonucleotides and brought out of the COY to a preheated BioRad T100 Thermal cycler. DNA annealing was performed by heating at 95 °C for 5 minutes followed by cooling to 25 °C at a rate of 0.1 °C/s, after which the tray and cover were immediately brought back into the COY chamber. The DNA was concentrated using 30 kDa MWCO centrifugal concentrators. The double stranded DNA was concentrated to 160 μM and buffer exchanged into high salt buffer (100 mM Tris HCl pH 6.8, 300 mM NaCl, 10% glycerol) and frozen anaerobically in liquid nitrogen, before being transferred to CHESS for the experiment.

### Small-angle X-ray Scattering

Anoxic SAXS (anSAXS) was performed at the ID7A station at the Cornell High Energy Synchrotron Source (CHESS) over multiple runs. The first experiment, where the oxygen dependence of the dimer-to-monomer transition was probed, used a 9.9 keV 250 μm x 250 μm X-ray beam with a flux of 7.1×10^11^ photons s^-1^. The second experiment, involving the binding of dimeric FNR to DNA, used a 9.9 keV 250 μm x 250 μm X-ray beam with a flux of 1.2×10^12^ photons s^-1^. For both runs, images were collected on an Eiger 4M detector, covering a range of *q* ≈ 0.009–0.5 Å^-1^. Each sample was centrifuged at 18,000 rcf for 8-10 min at a fixed temperature of 23 °C to remove aggregation immediately before loading onto a Superdex 75 Increase 10/300 column contained within the anSAXS chamber preequilibrated with elution buffer (100 mM Tris HCl pH 6.8, 10% glycerol, 150 mM NaCl). The sample was eluted at a flow rate of 0.5 mL/min, and the X-ray sample cell was set to a temperature of 23 °C. Roughly 800-1000 3-second exposures were collected per sample. Initial data processing and deconvolution were performed in BioXTAS RAW^42^.

To prepare samples of the FNR-DNA complex, *E. coli* FNR with an Fe content of 2.43 ± 0.72 per monomer was prepared anaerobically at 6.5 mg/ml (240 μM dimer concentration). A double-stranded 54-bp oligonucleotide containing the *nrdDG* promoter sequence was prepared at 160 μM in high salt buffer (100 mM Tris HCl pH 6.8, 300 mM NaCl, 10% glycerol). A sample with excess FNR was prepared by combining 300 μL of 120 μM FNR dimer with 100 μL of 160 μM oligonucleotide and concentrating to 100 μL with a 50 kDa molecular weight cutoff concentrator to remove any unbound DNA (MW 33 kDa) from the solution, with final concentration of 360 μM FNR and 160 μM concentration of oligonucleotide.

The final analysis was performed using the MATLAB implementation of EFA^15^ and REGALS^16^. After binning the datasets in both *q* and frames, SVD was performed to determine the number of significant singular values. For each dataset, the approximate start and end points for each component were established via preliminary evolving factor analysis (EFA)^15^. For each REGALS analysis, a background component concentration range was allowed to cover the entire range of the data using simple parameterization, while protein components were modeled with either simple or real-space parameterization (in the latter case using *D*_*max*_ from preliminary pair distance distribution analysis). Once component profiles were extracted, the forward scattering intensity *I*(0) and radius of gyration *R*_*g*_ were estimated by Guinier analysis, pair distance distribution analysis was done in GNOM^49^, and model fitting was performed in CRYSOL^50^ using 50 spherical harmonics. The model of *E. coli* FNR was obtained from the AlphaFold^51^ Protein Structure Database. Model building and geometry minimization of the FNR-DNA complex was performed in PyMol and Phenix^52^. Errors associated with *R*_*g*_ values are curve-fitting uncertainties from Guinier analysis.

## Data Availability

Requests for SAXS data and models can be sent to.

## Acknowledgments

The authors are grateful to Prof. Patricia Kiley and Dr. Erin Mettern at Univ. of Wisc-Madison for providing us with *E. coli* strains transformed with FNR plasmids and advice. We are also grateful to our first users, Drs. Amanda Byer and Max Watkins, for providing helpful feedback on the design and improvement of the anSAXS setup. Finally, we thank Dr. Steve Meisburger for suggesting FNR and Drs. Audrey Burnim, Amanda Byer, and Max Watkins for tips on protein expression and purification.

## Funding and additional information

SAXS was conducted at the Center for High Energy X-ray Sciences (CHEXS), which is supported by the National Science Foundation (BIO, ENG, and MPS Directorates) under award DMR-1829070, and the Macromolecular Diffraction at CHESS (MacCHESS) facility, which is supported by award 1-P30-GM124166-01A1 from the National Institute of General Medical Sciences (NIGMS), National Institutes of Health (NIH), and by New York State’s Empire State Development Corporation (NYSTAR). This work was supported by an NSF Graduate Research Fellowship under grant DGE-1650441 (to G. I.), NIH grant GM124847 (to N.A.) and startup funds from Cornell University (to N.A.). The content is solely the responsibility of the authors and does not necessarily represent the official views of the National Institutes of Health.

