## Supplemental Information for "Development of in-line anoxic small-angle X-ray scattering and structural characterization of an oxygen-sensing transcriptional regulator"

**This file includes:**

Figures S1-S7

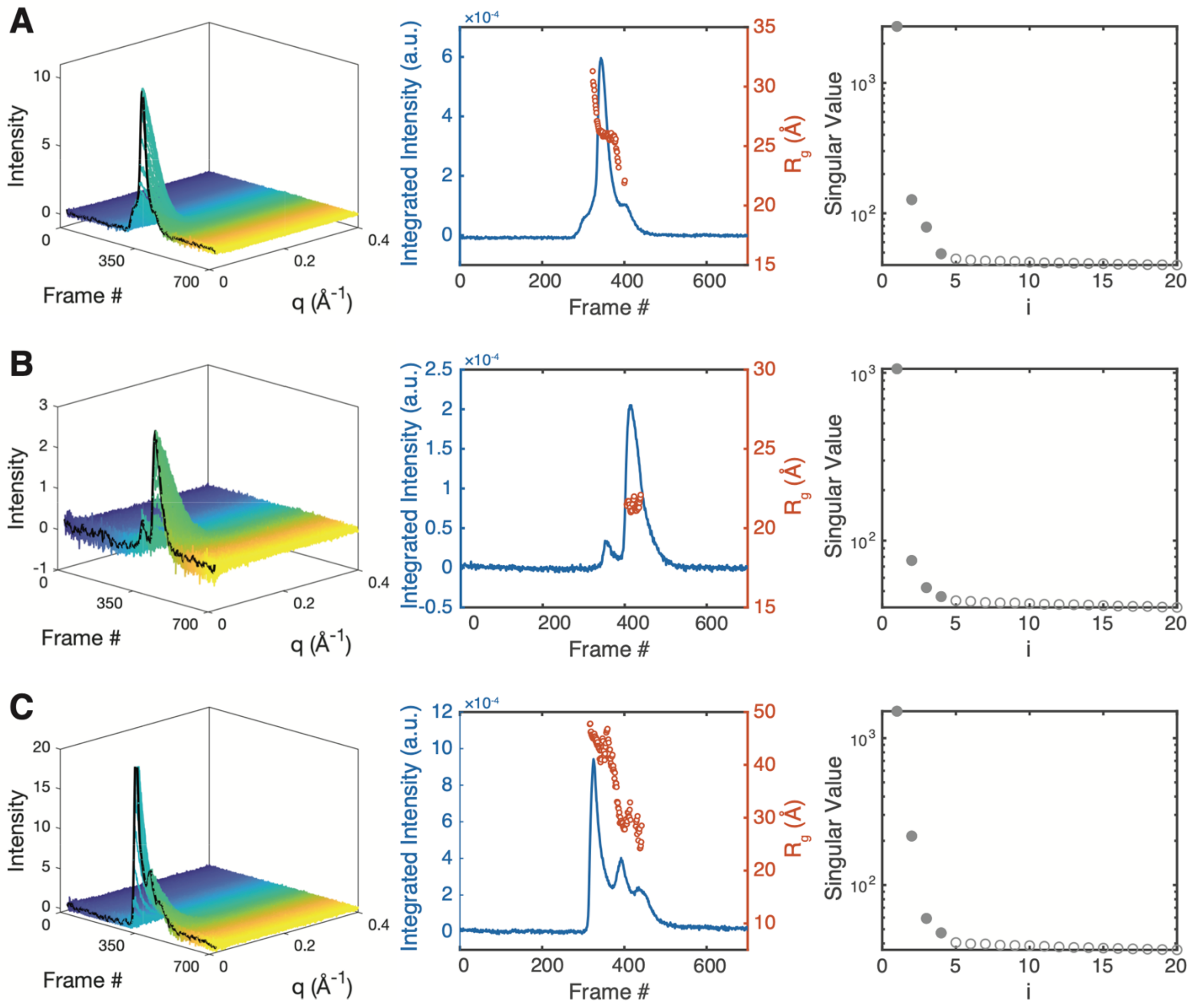


**Figure S1. Inspection of SEC-anSAXS datasets from *E. coli* FNR.** All samples were in 100 mM Tris HCl pH 6.8, 150 mM NaCl, 10 % glycerol; all FNR concentrations are dimer concentrations. **(A)** Left: SEC-anSAXS dataset of 164 µM FNR with an Fe content 3.27 ± 0.48 per monomer. Middle: The integrated intensities (blue) and $R_g$ values (red) from automated Guinier analysis reveal the elution of a large minor species, a main species with an $R_g$ ~26 Å (consistent with the value expected for dimeric FNR), and a small minor species. Right: SVD analysis shows there are four significant singular values within the dataset. **(B)** Left: SEC-anSAXS dataset of an aliquot of the sample described in panel A that was exposed to air for 5 min prior to data collection (Fe/monomer content was 1.91 ± 0.48). Middle: The integrated intensities (blue) reveal two relatively well-separated peaks, with the predominant species having an $R_g$ ~22 Å (consistent with monomeric FNR). SVD analysis indicates four significant components. **(C)** Left: SEC-anSAXS dataset of 360 µM FNR with 160 µM oligonucleotide containing the *E. coli nrdDG* promoter sequence. Middle: Integrated intensities (blue) show three overlapping peaks corresponding with a steady decrease in $R_g$ value (red). Right: SVD produces four significant values.

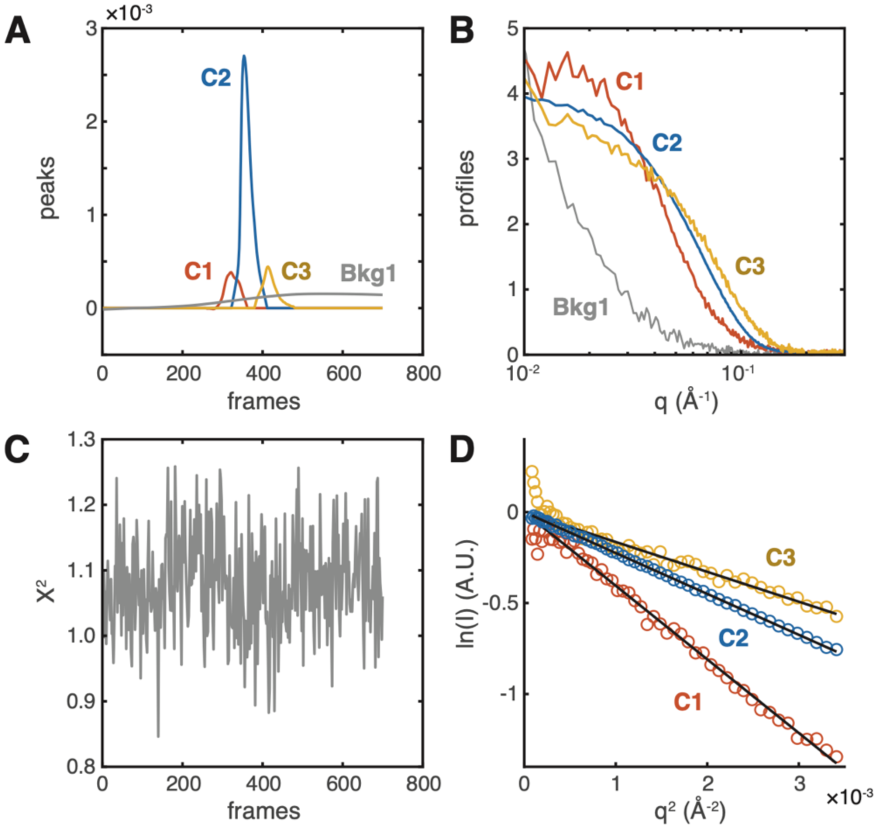


**Figure S2. REGALS analysis of anoxic FNR dataset.** REGALS deconvolution was performed with three protein components, C1 (orange, frames 258-362), C2 (blue, frames 322-410), and C3 (yellow, frames 380-480), and one background component, Bkg1 (grey), describing the change in background scattering over the entire elution. Simple parametrization was used for components with smoothing parameters set to [1e11, 1e1, 1e11, 1e14] for [C1, C2, C3, Bkg1]. **(A)** Extracted concentration profiles are consistent with the sequential elution of three species (red, blue, orange) over a changing background (grey), likely caused by material sticking to the X-ray cell over the course of the experiment. **(B)** Extracted profiles for the four components. **(C)** $\chi^2$ values are ~1 across all frames, indicating good agreement between the data and REGALS model. **(D)** Guinier analysis of protein scattering profiles in panel B produces $R_g$ values of 34.8 ± 0.5, 26.0 ± 0.1, and 22.2 ± 0.2 Å for C1, C2, and C3, respectively. These values are consistent with the elution of an aggregate species, FNR dimer, and FNR monomer.

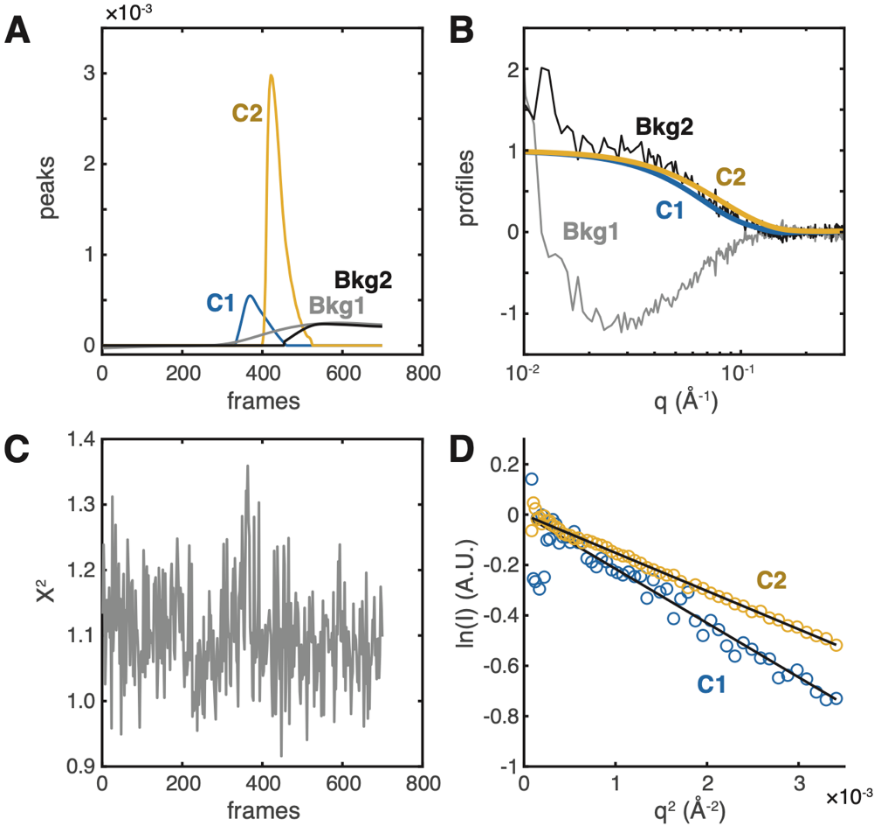


**Figure S3. REGALS analysis of oxygen-exposed FNR dataset.** REGALS analysis was performed using two protein components, C1 (blue, frames 332-460) and C2 (orange, frames 400-525) parametrized in real-space with $D_{max}$ of 100 Å and 110 Å respectively, and two background components, Bkg1 (grey, all frames) and Bkg2 (black, frames 454-endpoint), both with simple parameterization. Smoothing parameters were set to [1e12, 1e4, 1e14, 1e14] for [C1, C2, Bkg1, Bkg2]. **(A)** Extracted concentration profiles show a minor large component followed by the elution of a predominant smaller component. In this experiment, two components are required to describe the background scattering (grey, black). **(B)** Extracted profiles for the two protein components (blue and orange) and two background components (grey and black). **(C)** $\chi^2$ values are close to ~1 across all frames, indicating that the REGALS model is in good agreement with the data. **(D)** Guinier analysis of protein scattering profiles in panel B produces $R_g$ values of 25.4 ± 0.6 and 21.3 ± 0.1 Å for C1 and C2, respectively. These values are consistent with a small population of FNR dimer and a large population of FNR monomer.

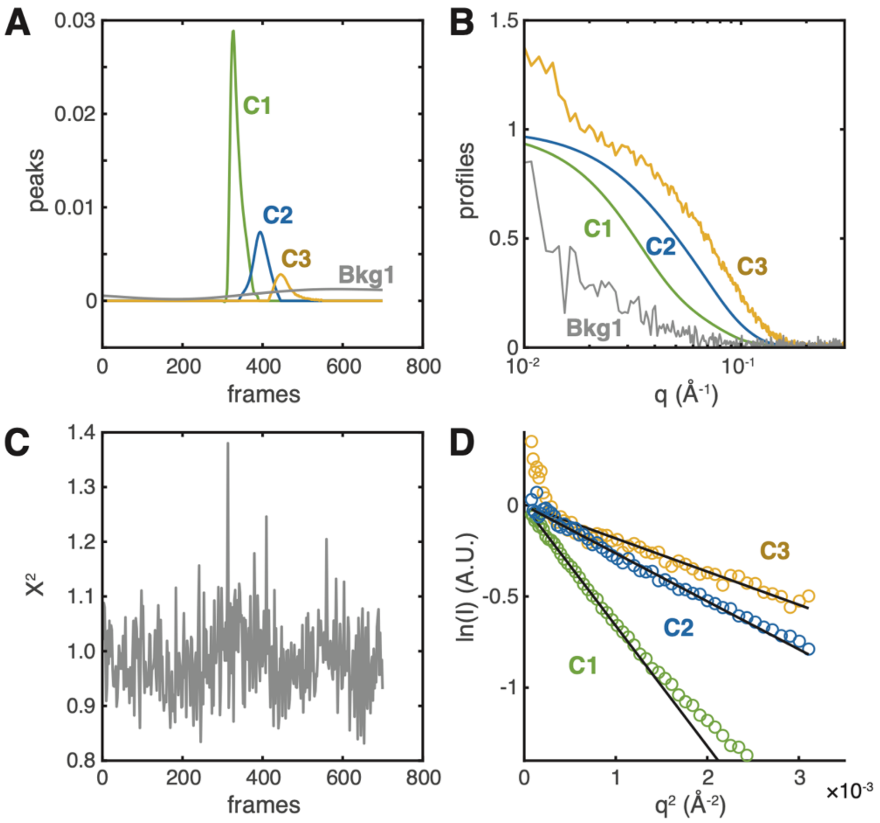


**Figure S4**. **REGALS analysis of DNA-FNR mixture dataset.** REGALS analysis was performed with three protein components, C1(green, frames 300-390, real-space parameterization with $D_{max}$ of 170 Å), C2 (blue, frames 340-445, real-space parameterization with $D_{max}$ of 115 Å), and C3 (yellow, frames 415-550, simple parameterization), and one background component Bkg1 (grey, simple parameterization). Smoothing parameters were set to [1e8, 1e10, 1e4, 1e14] for [C1, C2, C3, Bkg1]. **(A)** Extracted concentration profiles of three protein components show the sequential elution of three overlapping species (green, blue, orange) aver a changing background (grey), likely caused by material sticking to the X-ray cell over the course of the experiment. **(B)** Extracted profiles support the conclusion that there are three protein components (green, blue, yellow) and one background component (grey). **(C)** $\chi^2$ values remain close to ~1 throughout the frames indicating that REGALS analysis agrees well with data. **(D)** Guinier analysis of protein scattering profiles in panel B produces $R_g$ values of 44.5 ± 0.2, 28.1 ± 0.2, and 23.3 ± 0.3 Å for C1, C2, and C3, respectively.

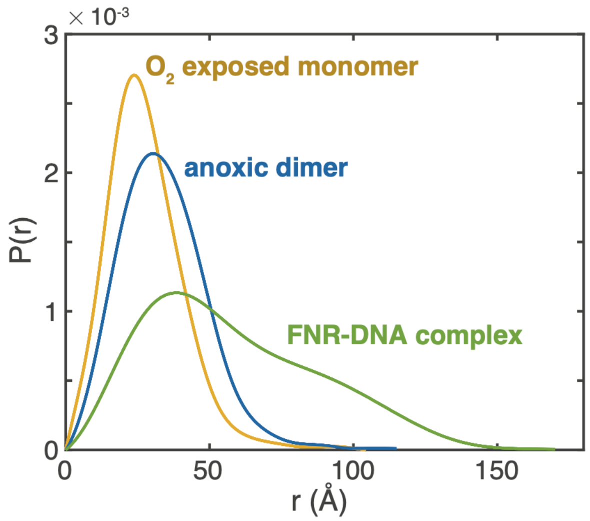


**Figure S5. Pair distance distribution analysis of dominant species in SEC-anSAXS datasets.** The pair distance distribution function, $P(r)$, of the *E. coli* FNR monomer produced by oxygen exposure (orange, component 2 in Figure S3) has a prominent peak, consistent with a globular shape. $P(r)$ of the FNR dimer (blue, component 2 in Figure S2) is largely monomodal, but the maximum dimension is slightly larger than that of the monomer, as expected. In contrast, $P(r)$ of the FNR-DNA complex (green, component 1 in Figure S4) has a very long dimension ($D_{max}$ of 170 Å) and displays a skewed shape with a peak at short distances (~30 Å), near the peak position of the dimer $P(r)$ and a shoulder at larger distances (~90 Å). This shape suggests that the FNR-DNA complex is overall an elongated species consisting of two copies of the FNR dimer held apart like a dumbbell.

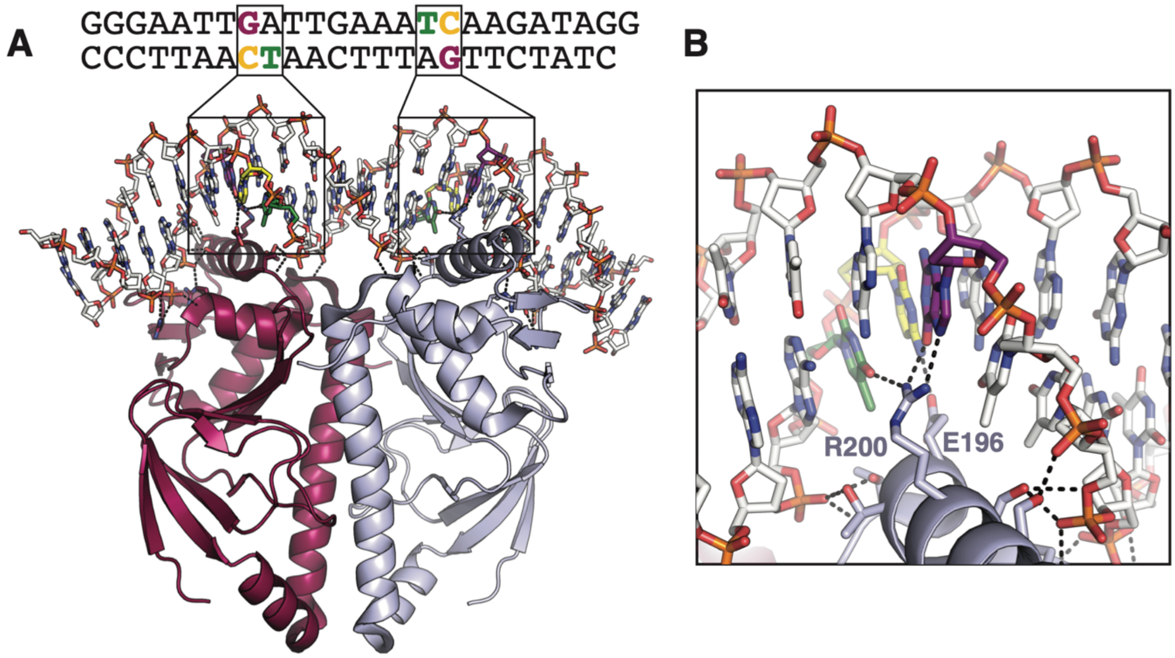


**Figure S6. Crystal structure of *B. japonicum* FixK2. (A)** The crystal structure of DNA-bound FixK2 dimer (PDB: 4i2o) shows that the DNA interacts with the protein both via its phosphate backbone and the nucleobases. Sequence-specific interactions are made by a triad formed by cytosine, thymine, and guanine bases (colored in DNA sequence and boxed in protein structure). **(B)** Close up of boxed region in panel A shows hydrogen-bonding interactions between FixK2 Arg200 and Glu196 with the cytosine, thymine, and guanine (CT/G) triad (colors matching the sequence in panel A).

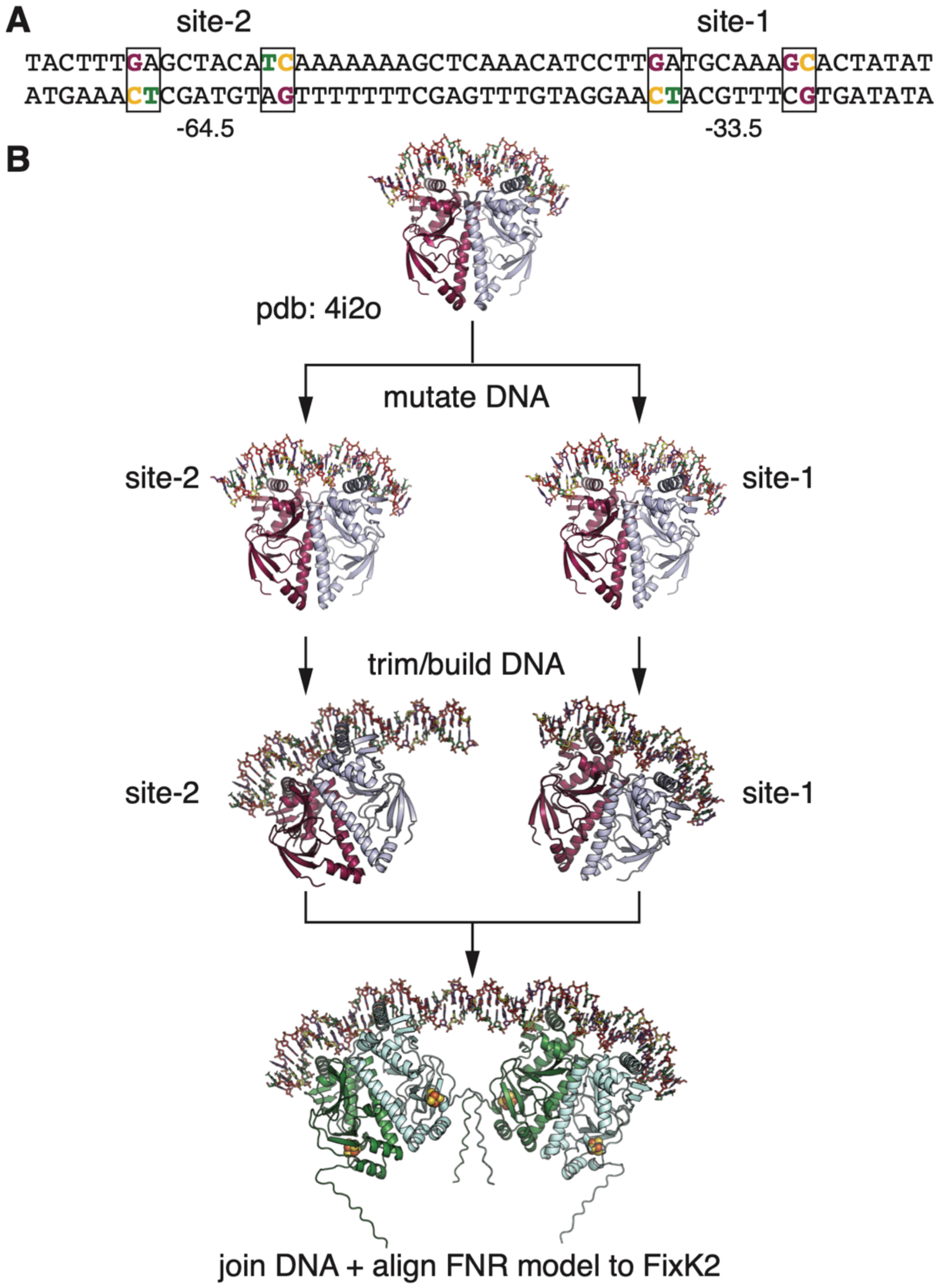


**Figure S7. Workflow to create model for *E. coli* FNR dimer bound to *nrdDG* promoter sequence. (A)** The *nrdDG* promoter sequence contains two FNR binding sites. The CT/G triads are predicted to hydrogen bond with FNR in a manner similar to that observed in Figure S6. In site-1, interactions with a conserved arginine in one of the FNR monomers are predicted to be made by two guanine bases rather than a guanine/thymine pair (as shown in Figure S6B). **(B)** The crystal structure of *B. japonicum* FixK2 (PDB: 4i2o) was used as a template to build a model of DNA-bound FNR. Two templates were produced, and the DNA sequence for each was mutated to match each of the binding sites in the *nrdDG* promoter sequence. After trimming extra nucleotides, the site-2 model was extended by B-form DNA with the intervening sequence, and site-1 was extended by an extra base pair. The two models were aligned by the overlapping base pair before removing redundant nucleotides and joining the chains. The AlphaFold2 model for *E. coli* FNR was aligned to each FixK2 dimer to produce a model of an FNR-DNA complex containing [4Fe-4S] clusters. The final model was manually refined and geometry minimized.
